# POME: Graph-based embeddings for partially observed mixed-type data

**DOI:** 10.64898/2026.09.10.750629

**Authors:** Fabian Woller, Lis Arend, Andreas M. Kist, Markus List, Farnaz Rahimi, Christel Sirocchi, David B. Blumenthal

## Abstract

Partially observed mixed-type (POM) data, as often encountered in clinical, epidemiological, and phenotypic datasets, are very common in biomedical research. Yet, advanced data analysis and machine learning based on POM data is complicated by their heterogeneous nature and often substantial fractions of missing values. While one promising way to overcome these issues is to compute vector-valued embeddings for POM datasets which can then be used for downstream analyses, existing embedding methods are mostly not designed for POM data. To address this gap, we developed POME (**p**artially **o**bserved **m**ixed-type data **e**mbeddings), a self-supervised model that yields low-dimensional representations of both samples and variables, using shared concept learning and a bipartite graph representation of the underlying POM data. We validated POME through extensive experiments on three real-world biomedical datasets, with diverse downstream tasks and objectives: POME achieves state-of-the-art imputation performance and produces high-quality patient representations that support not only unsupervised discovery of well-separated and clinically meaningful patients subgroups but also supervised predictive modeling and zero-shot representation space mining for use cases such as adjuvant therapy modality recommendation. POME is available as a Python package on GitHub (https://github.com/bionetslab/POME) and PyPI (https://pypi.org/project/pome-py).

## 1 Introduction

Clinical and phenotypic datasets in biomedical research often present analytical challenges due to their inherent heterogeneity in feature types. Additionally, experimental designs and data acquisition protocols frequently result in datasets with a significant proportion of missing values, which further complicates downstream analyses [1, 2]. Since many data analysis workflows are not designed to operate directly on incomplete or sparse data matrices, researchers must either employ deletion-based strategies such as complete-case analysis or pairwise missing value deletion, or apply one of numerous available data imputation techniques [3]. However, both deletion- and imputation-based strategies for handling missing values have been shown to introduce systematic bias under many realistic missingness mechanisms, potentially compromising the validity of downstream analyses [4, 5].

One promising way to overcome these issues is to compute vector-valued embeddings for such partially observed mixed-type (POM) datasets, which can then be used for downstream analyses. This approach has the advantage of not having to choose appropriate strategies for handling missing data or variable integration. Rather, the embedding model itself should be designed in a way such that resulting embeddings directly reflect similarities across variables of different types and by integrating the information hidden in the sparsity patterns of the dataset.

However, only very few existing embedding methods can handle POM data. Classical principal component analysis (PCA) is limited to numeric variables and cannot handle missing data [6], factor analysis of mixed data [7] supports mixed-type data but is still not applicable when missing data is present, and also the non-linear *t*-distributed stochastic neighbor embedding method (t-SNE) [8] does not integrate missing data into its core modeling. While t-SNE could be applied to partially observed data by computing the underlying distance matrix via a suitable missingness-aware distance measure (e. g., by using pairwise removal), extension to mixed-type data is less straightforward as no guidelines on how to combine variables of different data types exist. To the best of our knowledge, the only classical embedding method that fulfills all requirements of POM datasets (i. e., also handles categorical variables with more than two categories) is the uniform manifold approximation and projection method (UMAP) [9]: As t-SNE, UMAP does not integrate missing data into its core modeling, but can be applied to partially observed data through usage of a suitable distance measure. Furthermore, the authors provide a guideline on how to integrate variables of different types by computing the intersection of two separately fitted UMAP models [10].

Beyond classical approaches, scVI [11] is used in single-cell transcriptomics to embed numerical gene expression data while accounting for categorical confounding factors. Another popular multi-omics integration and embedding tool is MOFA [12, 13], which is capable of handling both numeric and ordinal variables, but is limited to binary categorical variables. There are also deep learning-based data imputation methods for POM data such as AutoComplete [14] or TabINR [15] that could be adapted to compute POM embeddings by extracting the models’ latent spaces. However, we are not aware of any deep learning model for POM data that comes with a ready-to-use implementation which would support such a usage.

To close this gap, we here present a new embedding method called POME (short for “**p**artially **o**bserved **m**ixed-type data **e**mbeddings”) specifically designed for POM data. POME is a self-supervised model that yields representations of samples, variables, and variable values. It uses shared concept learning to learn variable semantics based on a graph representation of the input data that enables the joint modeling of variables with heterogeneous data types and at the same time naturally encodes missing values as absent edges within the graph structure. POME’s setup is similar to classical approaches such as PCA, t-SNE, or UMAP, in the sense that it aims to capture the intrinsic structure of a specific dataset. Like PCA and UMAP, it is also capable of embedding samples unseen during training.

We tested POME on three biomedical datasets: The TCGA-LUAD dataset contains clinical and phenotype data from lung adenocarcinoma patients from a standardized and cleaned version of the TCGA Pan-Cancer cohort [16] (566 patients and 84 variables). Our second dataset HANCOCK [17] comprises ICD-10 codes, demographic annotations, as well as blood, pathological, and clinical outcome data (86 variables in total) for 763 head and neck cancer patients. Our third analyzed dataset is a subcohort of MIMIC-IV [18] curated by Rahimi et al. [19], with data for cancer patients treated with chemotherapy. It contains averaged values of 100 laboratory variables for 4,428 admissions over the 14 days prior to discharge, as well as binary aplasia and neutropenic fever labels that indicate if these chemotherapy side effects occurred within post-discharge observation windows.

Our results showed that POME produces versatile representations that are useful for several downstream tasks. First, we demonstrate that POME can be used for embedding-based data imputation, achieving performances comparable to state-of-the-art imputation methods. Second, we show that the learned embeddings are highly informative for unsupervised patient stratification, facilitating the identification and visualization of clinically meaningful patient clusters. Third, we find that POME’s embeddings substantially outperform UMAP in linear probing tasks, indicating superior utility for predictive modeling. Fourth, using the HANCOCK dataset, we demonstrate that POME’s embeddings can be leveraged for adjuvant therapy modality recommendation, where adherence to embedding-based recommendations is associated with improved patient survival. Finally, we conduct an exploratory analysis of POME’s variable representations on the MIMIC-IV dataset to investigate if POME can learn variable semantics that are plausible from a biomedical point of view.

## 2 Results

### 2.1 Overview of the POME model

In this section, we provide a high-level description of the POME model, further details can be found in Methods. POME is designed for any POM dataset of the form

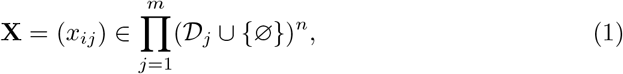

where *i* ∈ *S* = {1, …, *n*} are the saLJmples, *j* ∈ *V* = {1, …, *m*} are the variables, *D*_*j*_ is the domain of variable *j*, and 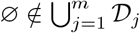 is a special symbol encoding missing data. Let *C* ⊆ *V* be the subset of categorical variables (binary or multiclass), *N* = *V* \ *C* be the subset of numeric variables, and ℬ be a set of *B* bins to discretize the domain *D*_*j*_ of a numeric variable *j* ∈ *N* via a suitable binning strategy *b*_*j*_ : *D*_*j*_ → ℬ. The binning functions *b*_*j*_ are required to represent numeric variables as nodes in POME’s graph-based model, in contrast to categorical variables, which are inherently discrete. The number of bins *B* is a hyperparameter and is the same for all numeric variables such that the set of bins ℬ is shared across all numeric variables. As default underlying all results presented in the main article, we used *z*-score binning with *B* = 15 bins (ablations showed that POME is robust with respect to both the type of the binning function and the number of bins *B*, see Fig. S1).

Moreover, let O = {(*i, j*) ∈ *S* × *V* | *x*_*ij*_≠ ∅} be the set of observed sample-variable pairs and let ℬ_*j*_ = {(*b, j*)|*b* ∈ ℬ} be a set of “bin nodes” for a fixed numeric variable *j* ∈ *N*. Then POME represents the dataset *X* as the following bipartite graph (Figure 1a):

**Fig. 1.**
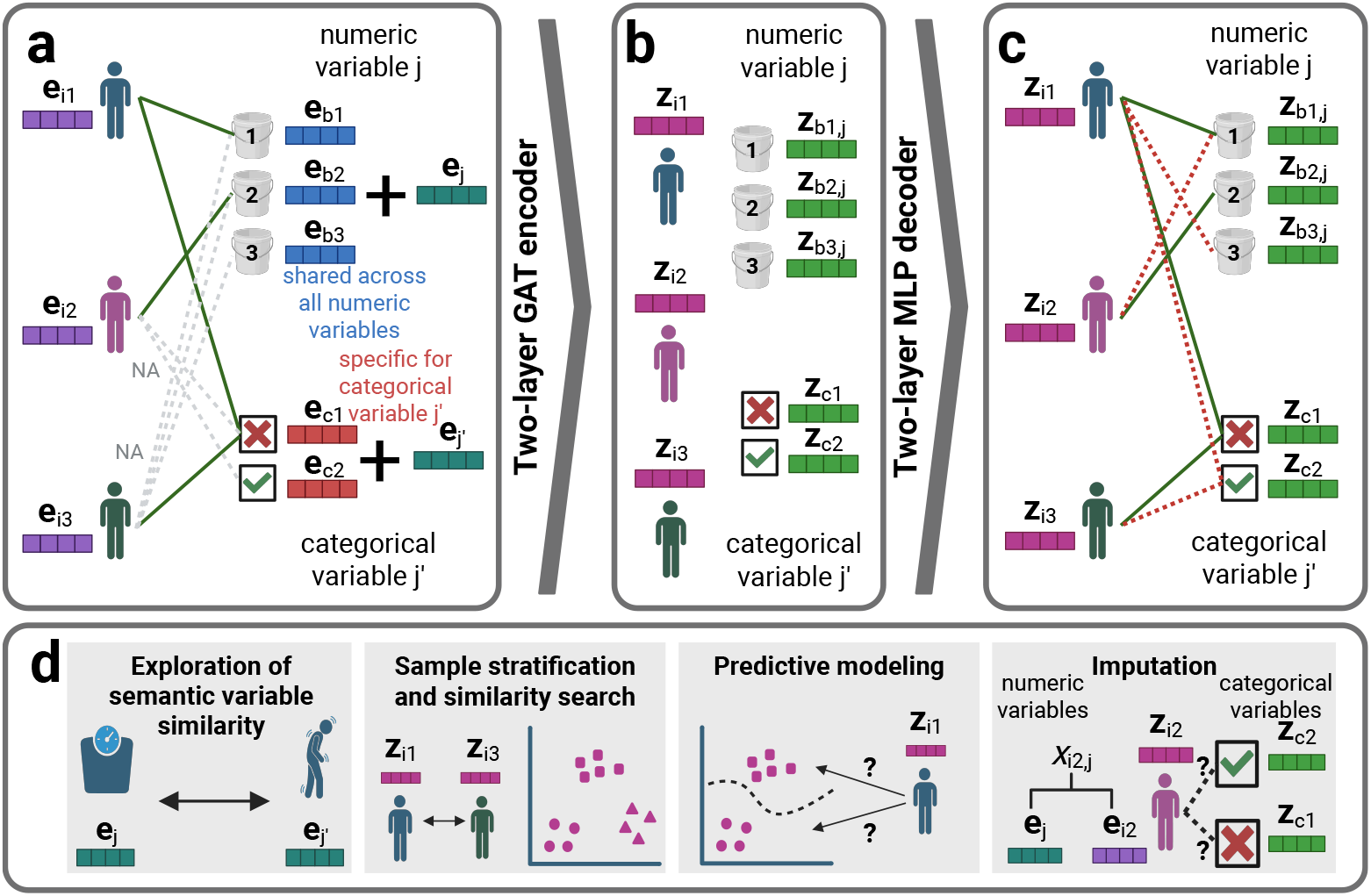
Overview of POME. **a** POME’s bipartite graph representation of POM data, here for simplicity with *B* = 3 bins for numeric variables. Sample embeddings are initialized with individual encodings (**e**_*i*1_, **e**_*i*2_, **e**_*i*3_), while variable value embeddings are initialized as the sum of a variable-specific encodings (**e**_*j*_, **e**_*j′*_) and a value-specific encodings. For numeric variables, value-specific bin encodings are shared across all variables (**e**_*b*1_, **e**_*b*2_, **e**_*b*3_). For categorical variables, value-specific category encodings (**e**_*c*1_, **e**_*c*2_) are created per variable individually. The node attributes are learned together with the model weights. **b** A GNN encoder ENC_***ϕ***_ (implemented as a two-layer graph attention network (GAT) [20]) learns embeddings for all samples (**z**_*i*1_, **z**_*i*2_, **z**_*i*3_), variable values (**z**_*c*1_, **z**_*c*2_) and bins (**z**_*b*1,*j*_, **z**_*b*2,*j*_, **z**_*b*3,*j*_). **c** A two-layer MLP decoder DEC_***θ***_ reconstructs the graph from the embedding space. In each training epoch, one non-edge is randomly sampled for each observed sample-variable pair (*i, j*) ∈ *O* (dotted red lines). **d** POME’s sample and variable representations can be used for various downstream tasks, including exploratory analysis of variable semantics, sample stratification and similarity search, predictive modeling, as well as imputation of missing values. Created in BioRender. Woller, F. (2026) https://BioRender.com/7s01yxh.

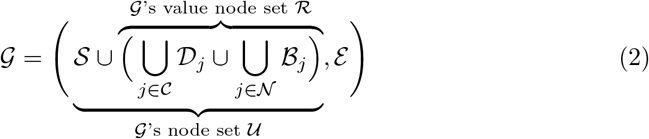

The intuition behind this graph representation is to match samples to their corresponding variable value, for each sample-variable pair. Hence, the edge set ℰ contains one sample-to-class edge (*i, x*_*i,j*_) ∈ *S* × *D*_*j*_ for each observed sample-variable pair (*i, j*) ∈ *O* with *j* ∈ *C* and one sample-to-bin edge (*i*, (*b*_*j*_(*x*_*i,j*_), *j*)) ∈ *S* × ℬ_*j*_ for each observed sample-variable pair (*i, j*) ∈ *O* with *j* ∈ *N*. This graph representation very naturally represents missingness: contains exactly one edge for each observed sample-variable pair; for non-observed pairs (missing values), the edge is simply omitted (Figure 1a). Note that this model assumes that categories of categorical variables are mutually exclusive, i. e., that there is only one correct category for each sample.

Using the bipartite graph representation *G*, POME trains a message-passing graph neural network (GNN) encoder

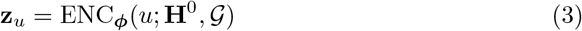

that produces node embeddings **z**_*u*_ ∈ ℝ^*d*^ of size *d* (hyperparameter) for all nodes *u* ∈ *U* of *G* (Figure 1b) and a link prediction decoder

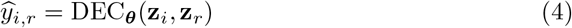

that yields edge scores 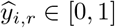 for all pairs (*i, r*) ∈ *S* ×ℛ of sample and value nodes (Figure 1c). 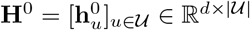 is the matrix of node embeddings used as inputs for the first layer of the GNN encoder. These initial node embeddings are defined as follows:

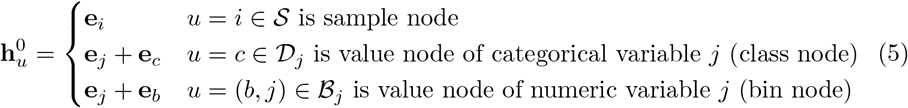

Here, **e**_*i*_, **e**_*j*_, **e**_*c*_, **e**_*b*_ ∈ ℝ ^*d*^ are sample, variable, class, and bin encodings that are learned together with the parameters ***ϕ*** and ***θ*** of the encoder and decoder (Figure 1a). The encoding **e**_*j*_ of a fixed variable *j* ∈ *V* is shared across all corresponding value nodes of *j*. The encoding **e**_*b*_ of a fixed bin *b* ∈ ℬ is shared across all the bin nodes (*b, j*) of all numeric variables *j* ∈ *N*, since this bin represents a comparable range ordering across all numeric variables. This shared concept learning approach allows POME to learn variable and bin semantics. Note that the class encodings **e**_*c*_ of categorical variables are not shared across variables, since classes of different categorical variables do not have a joint interpretation, in contrast to numeric bins, which are generated by the same type of binning function.

In total, POME hence has

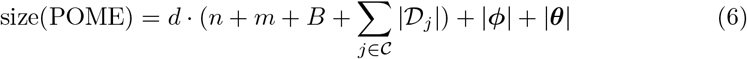

learnable parameters. We fit these parameters with a simple link prediction loss

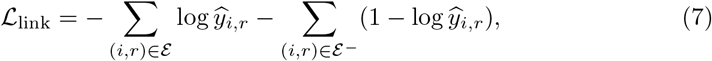

where ℰ^−^⊆ *S* × ℛ \ ℰ is a suitably sampled set of non-edges.

To support inductive reasoning, POME also allows to compute embeddings

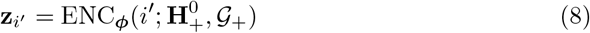

for new inference-time samples *i*^′^ ∈ *S*^′^ that are unseen during training (i. e., *S*^′^ ∩ *S* ≠ ∅). For this, each new sample is initially zero-encoded (yielding the augmented matrix of initial node embeddings 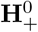), and a corresponding new node is added to the bipartite graph, with edges constructed based on *i*^′^’s POM data profile 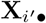 (yielding the augmented graph *G*_+_). Crucially, these edges are directed, pointing from value nodes to the new sample node *i*^′^. Like this, the new sample aggregates messages from its value nodes but never sends messages back. This leaves the representations of all training nodes (in particular the shared value nodes) identical to training, and ensures that all new samples can be embedded in a single forward pass through ENC_***ϕ***_ without perturbing the training graph or one another.

The trained POME model supports various downstream tasks (Figure 1d): The sample embeddings **z**_*i*_ can serve as input for unsupervised sample stratification, patient similarity search, or predictive modeling. The decoder DEC_***θ***_ can be used to impute data for non-observed sample-variable pairs (*i, j*) ∈ (*S* × *V*) \ *O*. And the learned variable encodings **e**_*j*_ enable exploratory analysis of variable semantics. In the following sections, we present the results of in-depth evaluations with respect to each of these downstream tasks.

### 2.2 Imputation of missing values

POME naturally supports imputation of values for unobserved variable pairs (*i, j*) ∈ (*S* × *V*) \ *O*. If the variable *j* is categorical, we impute the missing value *x*_*ij*_ as the highest-probability value

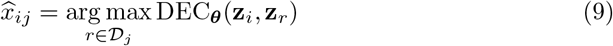

predicted by the decoder. For numeric variables, we equip POME with a lightweight regression head

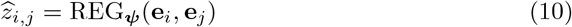

that predicts the *z*-score-transformed numeric values from the learned input encodings **e**_*i*_ and **e**_*j*_ of samples *i* ∈ *S* and numeric variables *j* ∈ *N*. We train REG_***ψ***_ with frozen input encodings **e**_*i*_ and **e**_*j*_ by minimizing the absolute error

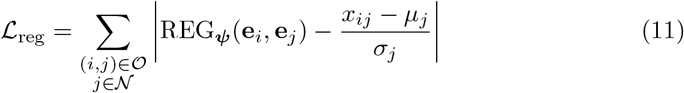

over all observed numeric entries, where *µ*_*j*_ and *σ*_*j*_ denote the mean and standard deviation of the observed values of the numeric variable *j* ∈ *N*. At inference time, we impute the missing values for unobserved sample-numeric variable pairs (*i, j*) ∈ (*S* × *N*) \ *O* by rescaling the predicted *z*-scores:

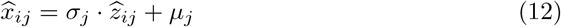

To benchmark POME’s imputation performance, we randomly placed missing values into the observed sample-variable pairs of the HANCOCK, MIMIC-IV, and TCGA-LUAD datasets, using simulated missingness ratios of 1%, 5%, 10%, 25%, and 50% and all variables (Table S1). For each missingness ratio and dataset, we generated 10 random missingness masks, fit POME with *O* subset to the unmasked sample-variable pairs, and quantified imputation performance by comparing *x*_*ij*_ and *x*_*ij*_ on the masked pairs, using multiclass accuracy for categorical variables and the range-normalized mean absolute error for numeric variables (each variable’s mean absolute error divided by its range in the unmasked data). We carried out the same experiments for three existing imputation methods that support mixed-type data: *k*-nearest-neighbor (*k*-NN) imputation [21], the tree-based imputation method MissForest [22], and the deep learning-based imputation method AutoComplete [14]. To obtain a condensed view on the results, for each simulated missingness mask, we ranked the four considered imputation methods according to the respective error measure for numeric and categorical variables. This way, we obtained averaged ranks per simulated missingness ratio across all three analyzed datasets.

The results are shown in Figure 2, with POME run with embedding size *d* = 64 (results for *d* ∈ {16, 32} are shown in Fig. S2–S3 and are slightly worse, indicate that POME’s imputation benefits from larger embedding sizes). Overall, we observe that POME clearly outperforms all competitors on categorical variables across all five simulated missingness ratios (Figure 2a) and is on par with MissForest on numeric variables, outperforming *k*-NN imputation and AutoComplete (Figure 2e). More specifically, on categorical variables, POME significantly outperforms all other methods on all datasets except for being on par with AutoComplete on MIMIC-IV (Figure 2b–d). On numeric variables, POME significantly outperfoms the best competitor MissForest on the HANCOCK and TCGA-LUAD datasets and is outperformed by it on MIMIC-IV (Figure 2f–h). These convincing results make POME an excellent go-to method for imputing missing values on POM data.

**Fig. 2.**
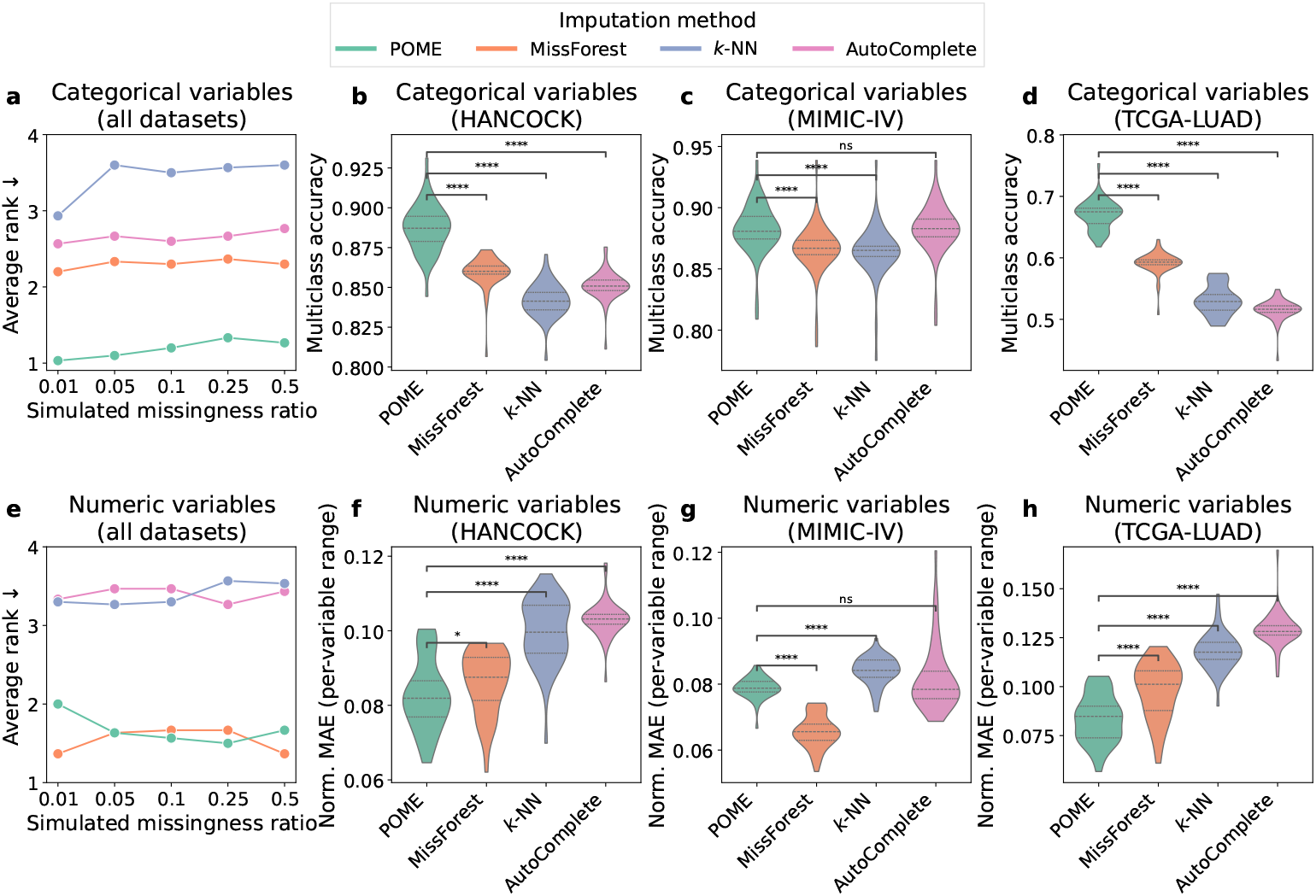
Results of imputation experiments with embedding size *d* = 64. **a** Average ranks of the compared methods on the categorical variables, jointly computed over the HANCOCK, MIMIC-IV, and TCGA-LUAD datasets (lower is better). **b–d** Multiclass accuracy scores on the categorical variables for each dataset separately, with violin plots visualizing results for all five simulated missingness ratios. **e** Average ranks of the compared methods on the numeric variables, jointly computed over the HANCOCK, MIMIC-IV, and TCGA-LUAD datasets (lower is better). **f–h** Mean absolute error on the numeric variables, normalized per variable by its min–max range, for each dataset separately, with violin plots visualizing results for all five simulated missingness ratios. Significance annotations are based on two-sided Mann-Whitney U tests.

### 2.3 Unsupervised sample stratification and visualization

One of the most important use cases of POME is that the sample embeddings **z**_*i*_ enable *de novo* identification of sample subgroups based on the joint information encoded in the heterogeneous set of variables of a POM dataset. To test POME with respect to this downstream task, we computed POME embeddings of sizes *d* = {16, 32, 64} using 10 different random seeds for the HANCOCK, MIMIC-IV, and TCGA-LUAD datasets (excluding targets and target-associated variables as specified in Tables S1– S2), clustered the embeddings via Leiden clustering [23] (which has the advantage of automatically deciding on the optimal number of clusters), and assessed clustering quality by computing Silhouette scores and Davies-Bouldin indices. We carried out the same workflow with embeddings computed with a customized version of UMAP which, as explained in the Introduction, is the only existing embedding method we are aware of that comes with published instructions on how to handle POM data.

Figure 3a–g shows the results. On two of the three datasets (TCGA-LUAD and MIMIC-IV), POME embeddings of size *d* ∈ {16, 32} yield substantially better Silhouette scores and Davis-Bouldin indices than UMAP embeddings of any size. Another interesting observation is that, on most datasets, the 16- and 32-dimensional POME embeddings tend to cluster better than the 64-dimensional embeddings. This is in line with a previous study on transductive graph-based representation learning, which also found that low-dimensional embeddings exhibit a better clustering behavior [24]. Furthermore, we observed that, compared to POME, UMAP embeddings yield clusterings with less variation in both clustering metrics. Since UMAP uses always the same *k*-NN graph representation of the underlying dataset as input, we hypothesize that this creates less variation in the structure of the final embeddings compared to the negative sampling approach integrated into POME.

**Fig. 3.**
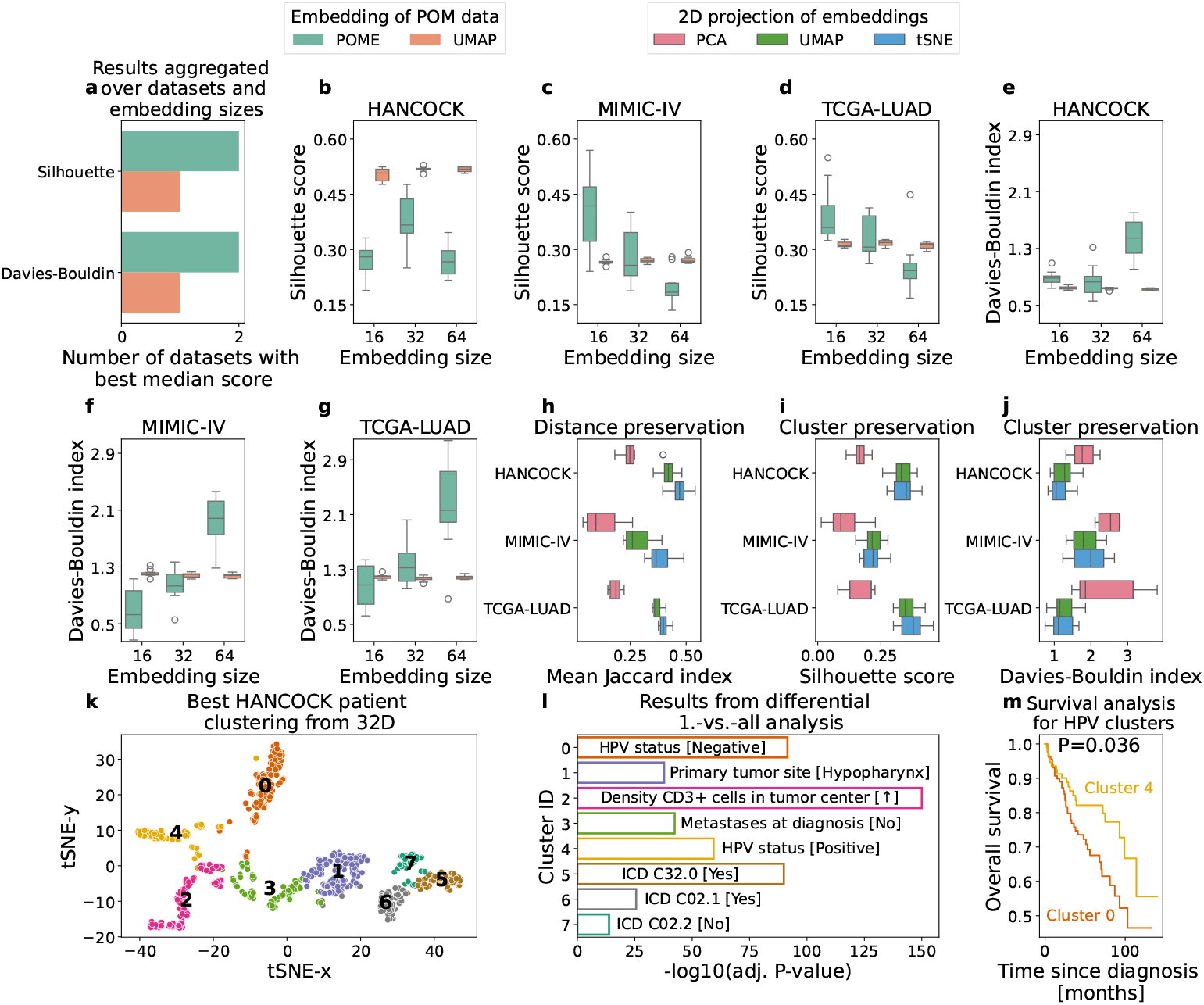
Results of unsupervised sample stratification and visualization experiments. **a** Numbers of datasets for which POME or UMAP embeddings lead to best median clustering scores across all tested embedding sizes *d* = {16, 32, 64}. **b–d** Silhouette scores of clusterings computed via Leiden clustering using POME and UMAP embeddings (higher is better). **e**–**g** Davies-Bouldin indices for the same clusterings as shown in (b)–(d) (lower is better). **h** Local neighborhood preservation between POME embeddings and 2D projections, aggregated across all embedding sizes *d* ∈ {16, 32, 64} (results for individual *d* in Fig. S4). **i, j** 2D preservation of clusterings computed in *d*-dimensional embedding spaces via Leiden clustering, quantified via Silhouette scores (i) and Davies-Bouldin indices (j) computed in 2D space. Results are aggregated across all embedding sizes *d* ∈ {16, 32, 64} (results for individual *d* in Fig. S4). **k** The t-SNE visualization of the 32-dimensional HANCOCK patient embedding with the highest Silhouette score in (b). Leiden clustering on the 32-dimensional embeddings led to eight patient clusters, which we use for coloring in two dimensions. **l** Top differential variables for the eight clusters shown in (k). The annotations in brackets after the variable names specific the within-cluster majority classes (categorical variables) or the direction of the fold change (numeric variable characteristic of cluster 2). **m** Survival analysis for the HPV-associated clusters 0 and 4.

Next, we asked which dimensionality reduction method is most appropriate to project POME’s *d*-dimensional sample embeddings to a two-dimensional (2D) Euclidean space for visualization purposes. We tested the three popular dimensionality reduction methods PCA, UMAP, and t-SNE. For each tested dimensionality reduction method *f* and each sample *i*, we quantified local distance preservation by computing the Jaccard index between the sets 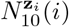 and 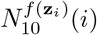 of 10 nearest neighbors of sample *i* retrieved in *d*-dimensional embedding space based on the embeddings **z**_*i*_ and in 2D Euclidean space based on the projections *f* (**z**_*i*_), respectively. Moreover, we quantified the preservation of the clusterings underlying the results presented in Figure 3b–g by computing Silhouette scores and Davies-Bouldin indices based on Euclidean distances of the embedding projections *f* (**z**_*i*_). As shown in Figure 3h–j, t-SNE projections performed best on all three datasets with respect to all three metrics. We thus recommend to visualize POME embeddings via t-SNE projections.

A t-SNE visualization of the clustering of POME embeddings for the HANCOCK dataset that corresponds to the best Silhouette score in Figure 3b is shown in Figure 3k. To identify characteristics of the eight patient clusters, we went back to the original POM data and performed one-cluster-versus-all differential analyses using missingness-aware versions of the Mann-Whitney U test or the *χ*^2^-test (depending on the variable datatype) for all clusters and all variables of the HANCOCK dataset. The characteristic variables with the lowest Bonferroni-corrected *P*-values are shown in Figure 3l, together with the most frequent category in the respective cluster (for categorical variables) or an indicator if values inside the cluster are on average higher than outside of the cluster (for cluster 2’s characteristic numeric variable). For all patient clusters except the negatively characterized cluster 7, the obtained characterizations are clinically plausible:

- Clusters 1, 5, and 6 are enriched with patients diagnosed with tumors at specific sites (the ICD-10 codes characterizing cluster 5 and 6 stand for malignant neoplasm of the glottis (C32.0) and of the border of tongue (C02.1), respectively).
- Cluster 3 is enriched with patients without metastases at the time of diagnosis.
- Cluster 2 is enriched with patients with immunologically active tumor centers with elevated densities of CD3-positive cells. In the original publication of the HANCOCK dataset, this has been identified as a positive prognostic marker [17].
- Clusters 0 and 4 are enriched with patients with negative and positive HPV infection status (note that since HPV status is measured only for patients that meet certain clinical criteria, HANCOCK’s “HPV status” variable has a third category encoding missing by design [17]). A positive HPV infection status is known as a positive prognostic marker in head and neck cancer [25] — a link which we could reproduce by comparing overall survival of patients grouped into the clusters 0 and 4 (Figure 3m, *P* = 0.036, log-rank test).

Overall, these results highlight the value of POME’s sample embeddings for unsupervised sample stratification: The embeddings yield clusterings with pronounced structure when used as input for standard Leiden clustering, can be faithfully visualized via t-SNE projections, and allowed us to rediscover clinically plausible and survival-associated patient subgroups in an unsupervised manner.

### 2.4 Adjuvant therapy modality recommendation

Another particularly promising potential use case for POME’s patient embeddings is to use them for patient similarity queries which could inform medical decisions such as selection of adjuvant therapy (AT) strategies in the context of tumor boards. To exemplify the value of POME in such scenarios, we carried out a retrospective AT recommendation study with the HANCOCK dataset, which contains information regarding AT modality and 5-year overall survival (OS) for patients diagnosed with head and neck cancer between 2005 and 2019 (Figure 4a, b). Specifically, we computed POME embeddings for all patients in the dataset (excluding targets, target-associated, and AT-associated variables as specified in Tables S1–S2), treated patients diagnosed in 2019 as “query patients”, and, for each query patient *i*, retrieved the set *N*_10_(*i*) of *k* = 10 patients diagnosed in earlier years whose embeddings are most similar to the embedding of *i*. We grouped *N*_10_(*i*) by AT modality, computed 5-year survival fractions for all subgroups of *N*_10_(*i*), retrospectively suggested the AT modality with the largest OS fraction as optimal AT modality for *i*, and assessed whether *i*’s actual AT modality matches the POME-based recommendation (if this is the case, we call *i*’s AT modality “POME-consistent”). Then, we computed 5-year OS fractions for query patients with and without POME-consistent AT modalities. We carried out this workflow 3 × 10 times, using embedding sizes *d* ∈ {16, 32, 64} and 10 POME embeddings per embedding size that were computed with different random seeds.

**Fig. 4.**
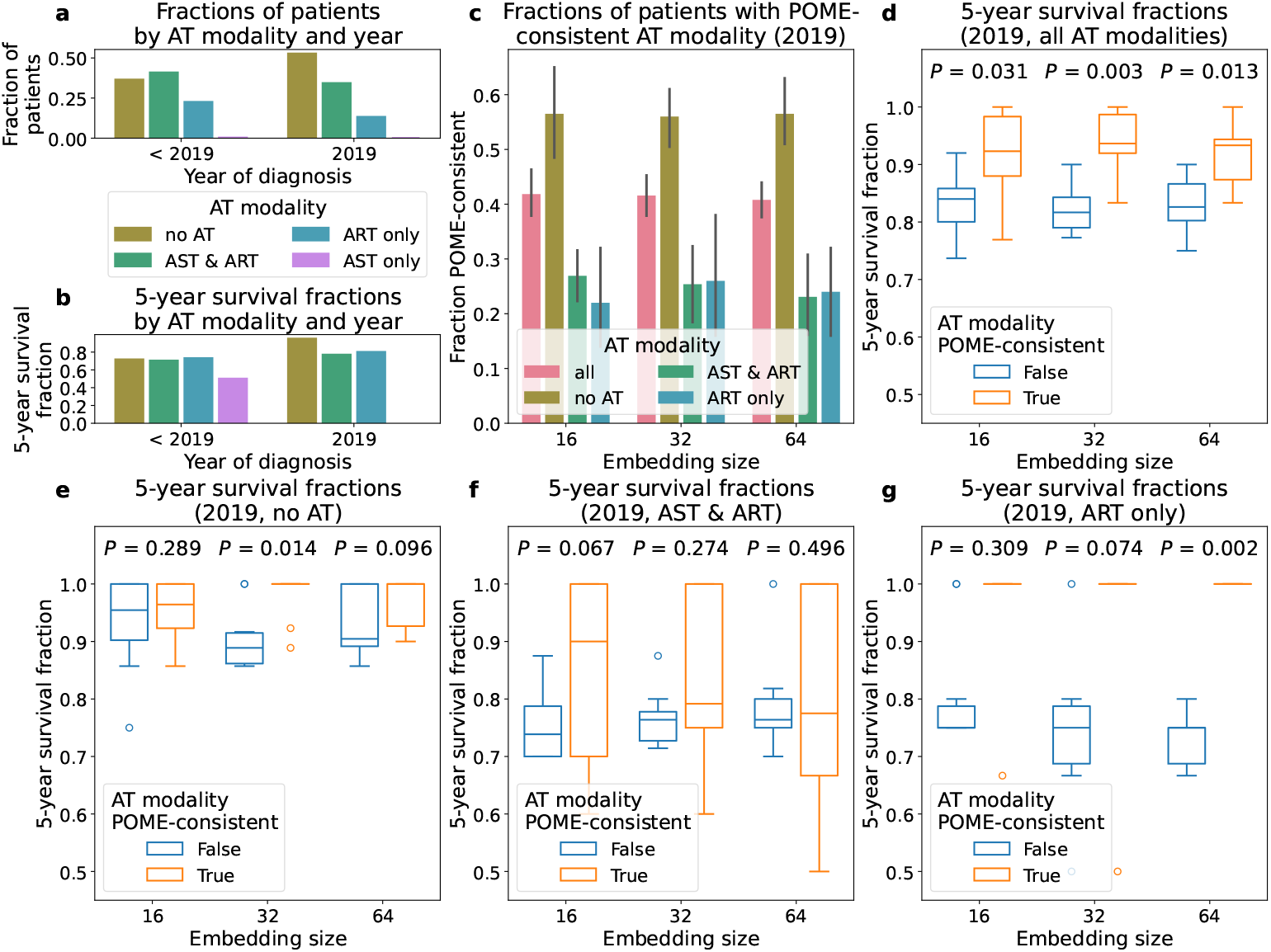
POME-based retrospective adjuvant therapy (AT) recommendation for the HANCOCK dataset. **a** Fractions of patients from the four AT modality groups “no AT”, “adjuvant systemic therapy and adjuvant radiotherapy” (AST & ART), “only adjuvant radiotherapy” (ART only), and “only adjuvant systemic therapy” (AST only), separately for the “query patients” diagnosed in 2019 (38 patients) and patients with earlier diagnoses (725 patients). Note that none of the query patients fall into the AST-only group. **b** 5-year survival fractions for the same patient groups as in (a). **c** Fractions of query patients with POME-consistent AT modality. The barplots show means over 10 independent POME runs, error bars depict 95% confidence intervals. **d** 5-year survival fractions of all query patients with and without POME-consistent AT modalities. **e–g** 5-year survival fractions of query patients with and without POME-consistent AT modalities, separately for each of the three AT modality groups containing query patients. *P*-values are computed with the one-sided paired Wilcoxon signed-rank test (alternative hypothesis: 5-year survival fractions are larger for query patients with POME-consistent AT modality).

The fractions of query patients with POME-consistent AT modalities are visualized in Figure 4c, the distributions of OS fractions for the query patients are shown in Figure 4d–g. For all the embedding sizes, OS fractions computed across all AT modality groups are significantly larger for query patients with POME-consistent AT modality (Figure 4d). *P*-values increase when computing separate OS fractions for the three AT modality groups present among the query patients, but we still observe consistently larger OS fractions for patients with POME-consistent AT modalities across embedding sizes and AT modalities (Figure 4e–g, median OS fractions are larger for patients with POME-consistent AT modalities in all 9 comparisons). A particularly interesting finding is that, when using POME embeddings of size *d* = 32 to determine POME-consistency, OS fraction differences are statistically significant for the subgroup of query patients who did not receive any AT (Figure 4e). This subgroup shows the largest OS fraction in the cohort of query patients (Figure 4b), and for most of them, POME suggests “no AT” as the most appropriate AT modality (Figure 4c). Interestingly, the OS fraction drops when this is not the case (Figure 4e), indicating that our POME-based AT modality recommendation workflow correctly identified patients in the HANCOCK dataset who did not but should have received AT.

### 2.5 Linear probing

Next, we evaluated POME’s patient embeddings via linear probing to assess whether POME’s embeddings capture information that is relevant for the prediction of held-out target variables. To analyze this, we used eight binary target variables from the HANCOCK, MIMIC-IV, and TCGA-LUAD datasets (Table S2). Based on the remaining variables, we first generated ten random 80-20 train-test sample splits and computed POME and UMAP embeddings of sizes *d* = {16, 32, 64} using five different random seeds, excluding variables directly associated with the targets (Table S2). Embeddings for test samples were computed using POME’s and UMAP’s inductive mode, which only fits the corresponding model on training samples and then generates embeddings for unseen samples. For POME, the number of training epochs was selected via an unsupervised geometric criterion of the embedding matrix in order to avoid overfitting (see Methods and Fig. S5). UMAP’s inductive mode had to be adapted in order to be applicable in the setting of combining two models for different data types (see Methods). We then fit logistic regression models only on training sample embeddings, and evaluated their performances on embeddings of unseen test samples. Since some target variables contained missing values, we omitted samples with missing target information in this analysis.

The results are shown in Figure 5, with predictive performances quantified as average precision (AP). We obtained better AP scores for seven out of eight target variables with POME than with UMAP embeddings (Figure 5a). The embedding size has a moderate effect on the predictive performances of logistic regression models trained on POME embeddings, with larger embedding sizes leading to better AP scores for most targets. Independently of the embedding method, the AP scores are consistently above the positive class ratios *p* of the binary target variables (dashed lines in Figure 5b–i), representing the AP score of a random baseline classifier that predicts the positive class with probability *p*.

**Fig. 5.**
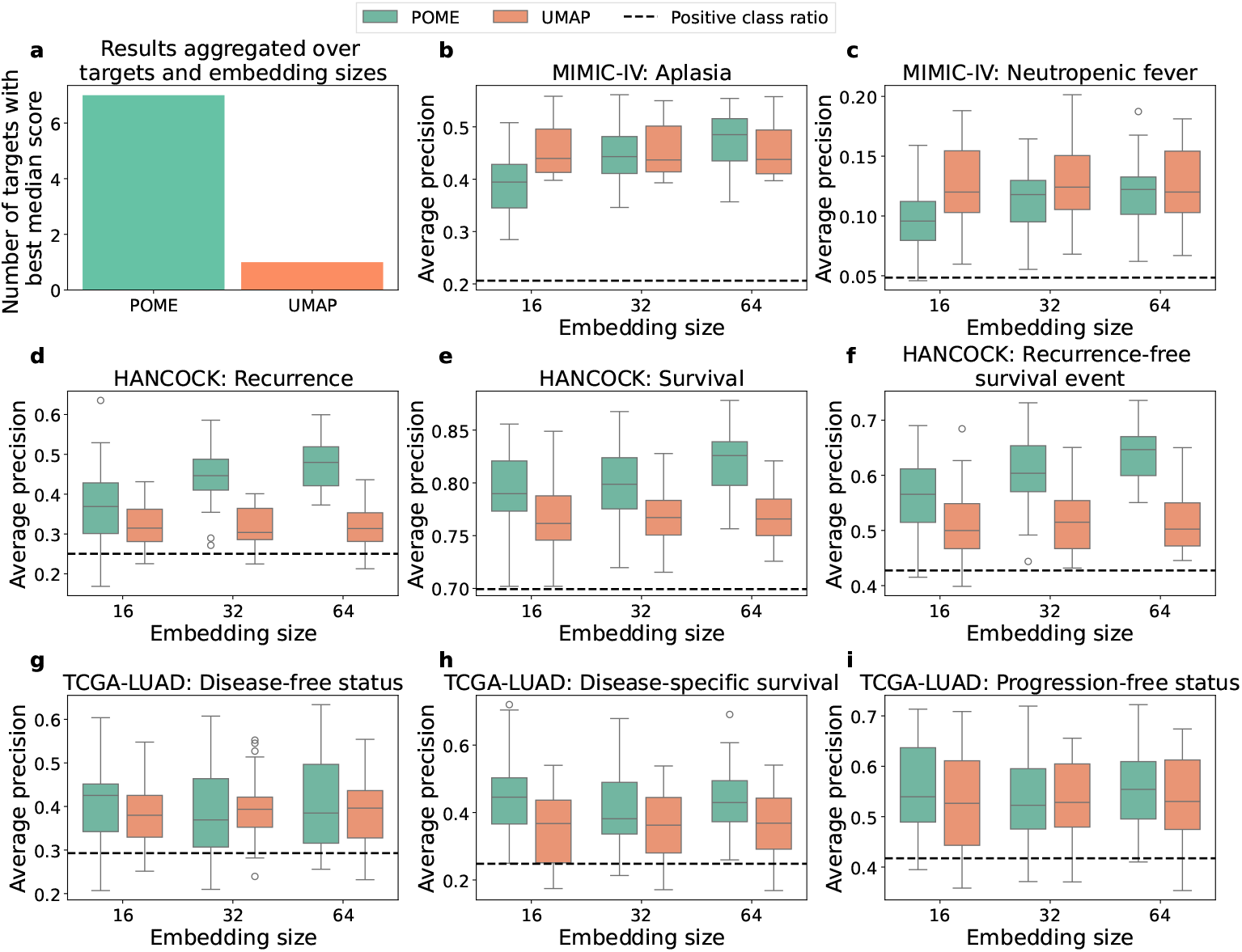
Results of linear probing experiments. **a** Numbers of target variables on the three datasets for which POME or UMAP embeddings lead to best median performance scores across all tested embedding sizes *d* = {16, 32, 64}. **b–i** Results for binary target variables, reported as average precision (AP). All box plots show AP scores of a logistic regression model fitted on training sample embeddings, and evaluated on unseen test sample embeddings. Embeddings of test samples are computed using POME’s and UMAP’s inductive embedding computation (that is, test samples were not used for fitting the POME and UMAP models). The evaluation includes ten training/test splits and five embeddings per embedding size generated with different random seeds (that is, each box plot summarizes 10 · 5 = 50 values).

Another observation is that UMAP embeddings generally yielded better results on the MIMIC-IV dataset, compared to the other two datasets. We hypothesize that UMAP has a slight advantage here because (without the two target variables) MIMIC-IV only consists of numeric variables, which means that only one “classical” UMAP model needs to be fitted. This is corroborated by the fact that logistic regression models trained on POME embeddings benefit from jointly representing numeric and categorical variables on a subset of target variables in the HANCOCK and TCGA-LUAD datasets, whereas models trained on UMAP embeddings never benefit from a joint representation (Fig. S6). The linear probing experiments hence corroborate the finding that POME yields information-dense sample representations.

### 2.6 Exploratory analysis of learned variable semantics

Finally, we investigated whether POME’s shared concept learning approach can capture meaningful variable semantics. We used the MIMIC-IV dataset for this purpose, fit POME on all variables including the two targets aplasia and neutropenic fever (*d* = {16, 32, 64}, 10 random seeds for each *d*), and retrieved the learned variable encodings **e**_*j*_ for all variables *j*. Moreover, we obtained feature importance scores from a previous study [19] where CatBoost [26] (and other) models were trained to predict aplasia and neutropenic fever from a temporal version of the MIMIC-IV dataset (as detailed in Methods, we aggregated the MIMIC-IV data during preprocessing to obtain the non-temporal version used for this study). Then, we computed cosine similarities between the target encodings and the encodings of all variables used as features by the CatBoost models, and computed Spearman correlation coefficients to assess if target-to-feature encoding similarity correlates with CatBoost feature importance.

As shown in Figure 6a, we obtained positive correlation coefficients for the aplasia target across all tested encoding sizes, but not for the neutropenic fever target. This is in line with the results of the linear probing experiment presented above, where POME’s sample embeddings yielded better results for aplasia than for neutropenic fever prediction (Figure 5b, c). To further understand if POME’s learned semantic of the aplasia variable is meaningful from a biomedical point of view, we clustered the 32-dimensional encodings that resulted in the largest Spearman correlation coefficient using Leiden clustering (t-SNE projection shown in Figure 6b) and then zoomed-in on the aplasia cluster (Figure 6c). Interestingly, the aplasia cluster contains many white blood cell-related variables such as monocytes and neutrophils in the vicinity of the aplasia variable, which are known indicators of immunosuppression [19].

**Fig. 6.**
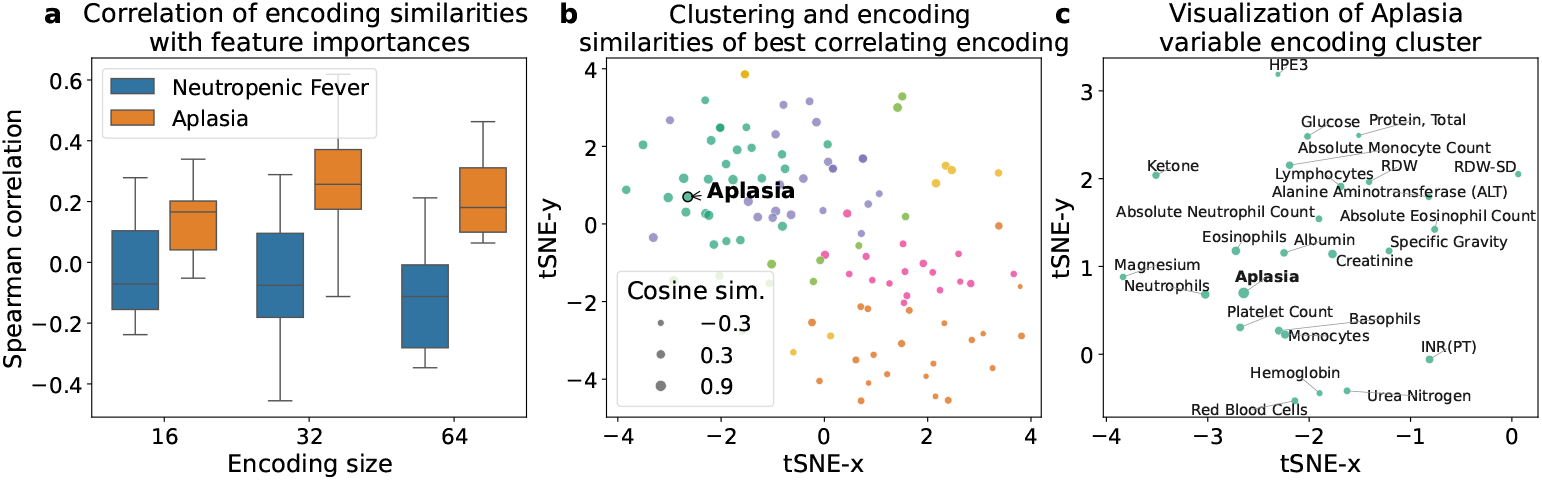
Results of exploratory analysis of variable semantics on the MIMIC-IV dataset. **a** Spearman correlation coefficients between (i) CatBoost feature importance scores from [19] for the target variables aplasia and neutropenic fever and (ii) cosine similarities between learned POME encodings of the targets and all other variables. **b** Leiden clustering of 32-dimensional variable encodings, visualized with t-SNE. **c** Detailed view on the cluster from (b) which contains the aplasia variable.

## 3 Discussion

In this article, we have presented POME, an embedding method explicitly designed for POM datasets. POME relies on a self-supervised transductive learning approach to compute both sample and variable value embeddings that capture the underlying structure of the input data. On three real-world biomedical datasets, we have shown that POME reaches state-of-the-art performance in diverse downstream tasks and objectives (imputation of missing values, unsupervised patient stratification, adjuvant therapy modality recommendation, linear probing, exploratory analysis of variable semantics). We also demonstrated that POME’s embeddings can be faithfully visualized by mapping them to 2D Euclidean space via standard t-SNE projections. Finally, POME is fast enough to be practically useful for real-world iterative POM data analysis workflows: on simulated data with 10,000 samples and 100 variables, fitting the model on standard GPU-powered hardware required less than two minutes (Fig. S7).

Despite these convincing results, we anticipate that it may be possible to further improve POME by increasing architectural complexity. Such modifications could include (i) adding (induced set) attention blocks over the learned initial sample and/or variable representations before the GAT encoder to further increase expressivity of POME’s sample and variable representations or (ii) replacing the simple MLP in the regression decoder REG_***ψ***_ by a transformer-based design to further boost imputation performance on numeric variables. For this work, we deliberately refrained from implementing such extensions as our primary objective was to assess the value of our key idea to leverage bipartite graph learning for POM data.

Moreover, it is important to note that POME’s current implementation is optimized for training time efficiency on GPU-powered hardware. While it is also usable on CPU architectures, training times increase substantially from less than one and two to more than three and eighteen minutes on simulated data with, respectively, 2000 and 10,000 samples (Fig. S7). Moreover, POME’s optimization for low training times comes at the cost of a substantial memory consumption: Both on GPU and on CPU hardware, fitting POME models for dataset with 10,000 samples and 100 variables required around 15 GB of main memory (Fig. S7). To ensure scalability to very large datasets, it could thus be beneficial to investigate computationally more efficient training strategies (e. g., randomly sampling a constant number of samples *i* ∈ *S* in each epoch).

In sum, POME is a versatile embedding method for POM data which relies on the key insight that partially observed discrete data can be represented very naturally as a non-complete bipartite graph between samples and variable values. POME learns representations of samples, variables, and variables values that can be used out-of-the-box for various downstream analyses in biomedical research and beyond. For many of these downstream tasks, we anticipate that it may be possible to further improve upon the results reported in this article through task-specific customization of POME; for some of them, we have sketched concrete strategies how this could be achieved. To facilitate such follow-up work and to ensure that POME is practically usable and accessible, we provide a thoroughly tested and extensively documented Python implementation that can easily be installed from PyPI: `pip install pome-py`.

## 4 Methods

### 4.1 POME: binning strategy

For discretizing the domain *D*_*j*_ of a given numeric variable *j* ∈ *N*, we use a *z*-score based binning function *b*_*j*_ : *D*_*j*_ → ℬ which is given by

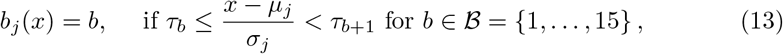

where ***τ*** = [−∞, 3.5, −3.0, …, −0.5, 0.5, …, 3.0, 3.5, ∞] and *µ*_*j*_ and *σ*_*j*_ denote mean and standard deviation over all non-missing values of variable *j*, respectively. Like this, bins also have a shared semantic interpretation across all numeric variables. This definition of *b*_*j*_ results in a number of bins *B* = 15. We also analyzed the effect of using fewer bins and an alternative non-linear discretization technique that yields smaller bin sizes with increasing value magnitudes (Fig. S1). However, we observed that *z*-score binning with *B* = 15 bins slightly outperforms these alternative binning strategies in terms of imputation performance, and we hence used this configuration for all results reported in the main article.

### 4.2 POME: encoder and decoder architectures

POME’s encoder ENC_***ϕ***_ consists of two GAT layers [20], with ReLU as a non-linear activation applied after the first layer. We use GAT layers because we hypothesize that different variables contribute non-uniformly to sample identity and thus want POME to be able to learn non-uniform edge weights used during message passing. We use two layers because, in a *k*-layer message-passing GNN, the number of distinct walks of length *k* between two nodes *u* and *u*^′^ in the underlying graph acts as a structural constraint on how much information can flow between *u* and *u*^′^ [27]. Our graph representation *G* contains exactly one walk of length *k* = 2 between samples *i, i*^′^ ∈ *S* for each variable *j* ∈ *V* such that (*i, j*), (*i*^′^, *j*) ∈ *O* and 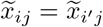, where 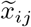 is the discretization of the original input data (i. e., 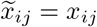 if *j* ∈ *C* and 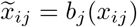 if *j* ∈ *N*). In our two-layer GAT encoder, information sharing between two sample nodes *i* and *i*^′^ is hence proportional to the missingness-aware Hamming similarity between their discretized data profiles. The output embeddings of the GAT are then normalized with respect to the *L*_2_ norm such that POME’s sample and variable embeddings **z**_*u*_ live on the unit sphere. Consequently, the cosine similarity 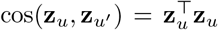 and the squared Euclidean distance ∥**z**_*u*_ − **z**_*u*_*′* ∥ ∝ (1 − cos(**z**_*u*_, **z**_*u*_*′*)) are appropriate similarity and distance measures for POME’s embedding space, and are thus used for all embedding similarity and distance computations in this article.

The decoder DEC_***θ***_ is a two-layer MLP with ReLU activation applied after the first layer, and uses concatenations [**z**_*i*_; **z**_*r*_] of sample and value node embeddings as input. After the second layer, a sigmoid activation converts the scalar outputs into edge probability values 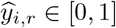. The regression decoder REG_***ψ***_ used for imputation of numeric variables is a three-layer MLP with two hidden layers of width 2*d*, each followed by a ReLU activation and 10% dropout, and a final linear layer producing a single scalar output. As input, it uses concatenations [**e**_*i*_; **e**_*j*_] of learned sample and variable encodings. The scalar output is the predicted *z*-score normalized value of the variable *j* for sample *i*, which is de-normalized during inference using *j*’s empirical mean and standard deviation estimated from the training data.

### 4.3 POME: inference-time adaptations for inductive reasoning

Since the initial encodings 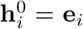 of training samples *i* are learned during training, we cannot use the same encoding strategy for new inference-time samples *i*^′^ unseen during training. Instead, we encode them via the null vector 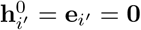. To prevent that this training-inference asymmetry leads to out-of-distribution embeddings **z**_*i*_*′* of inference-time samples *i*^′^, we remove the forward links (*i*^′^, *j*) ∈ *O* between *i*^′^ and its value nodes *j* in both message passing layers of POME’s trained GAT encoder and retain only the backward links (*j, i*^′^). That is, while the edges of the training graph are undirected, inference-time samples exclusively receive messages from their value nodes and never send messages back.

As a consequence, the latent representations 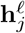 of all value nodes *j* are, in every layer ℓ, exactly identical to the ones obtained during training, i. e., the semantically meaningless zero encodings **e**_*i*_*′* cannot distort them at any stage of the forward pass. For the same reason, inference-time samples do not interfere with each other: **z**_*i*_*′* is fully determined by the observed values of *i*^′^ and independent of which other samples are embedded alongside it, so arbitrarily many inference-time samples can be embedded in a single forward pass through the frozen encoder. The only conceivable remaining influence of **e**_*i*_*′* is mediated by the self-loop that the GAT layers add to every node, including *i*^′^. In the first layer, this self-loop is in fact *not* down-weighted by the attention mechanism: because 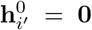, its unnormalized attention logit is exactly zero, such that its attention weight amounts to approximately the uniform share 1*/*(1 + |{*j* : (*j, i*^′^) ∈ *O*}|) – the same order of magnitude as the self-loop weights of the training sample nodes. It nevertheless propagates no information, since the message it carries, 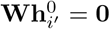, vanishes identically (its sole effect is that the attention softmax uniformly shrinks the weights of the value nodes). The first-layer representation 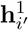 is hence a function of the observed values of *i*^′^ alone, and so is **z**_*i*_*′*, because the second-layer self-loop of *i*^′^ merely re-uses the already data-derived 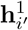.

### 4.4 POME: negative sampling and training policies

For the computation of the link prediction loss, we need to draw samples of non-existing (negative) edges ℰ^−^, since their number is significantly higher than the number of existing (positive) edges. In our implementation, we create balanced sets of positive and negative edges in each training epoch by randomly sampling one negative edge for each sample-variable pair that is also associated with a positive edge: for each sample-variable pair (*i, j*) ∈ *O*, we random_{_ ly draw one possible sam_}_ple-value pair from the set of all possible negative edges 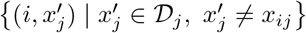 if *j* is categorical, or from the set {(*i*, (*b*^′^, *j*)) | (*b*^′^, *j*) ∈ ℬ_*j*_, *b*_*j*_(*x*_*ij*_) ≠ *b*^′^} if *j* is numeric. This implies that |ℰ|= |ℰ^−^| = |*O*| holds in each training epoch, and also ensures that no sample-value pairs of non-observed sample-variable pairs (*i, j*) ∉ *O* are (possibly erroneously) sampled into the set ℰ^−^ of negative edges.

For training our models, we use the Adam optimizer [28] with a learning rate of 0.01 and initialized the parameters via Xavier initialization [29]. We provide two training policies: one designed for transductive reasoning (no new samples arrive after training) and one for inductive reasoning (trained encoder ENC_***ϕ***_ should be used to embed inference-time samples unseen during training). For the transductive setting (used to generate results shown in Figures 2 to 4 and 6), we train POME for a fixed number of 2000 epochs, as in a transductive learning setting, we do not aim at generalization but want to obtain embeddings that best reflect the structure of the input data. This choice is justified by the fact that increasing the number of epochs led to improving imputation results (Fig. S8). In the transductive setting, POME can hence essentially be viewed as an anytime algorithm: The representational value of the embeddings increases with increasing number of training epochs, with diminishing returns the longer we train the model.

For the inductive setting (used to generate results shown in Figure 5), we determine the number of training epochs via a 3-fold cross-validation procedure over the training samples. All folds are trained in parallel for the same number of epochs, and every 10 epochs we inject each fold’s held-out validation samples inductively into its (frozen) training graph. We then compare the sets of embeddings of training and validation samples with a label-free geometric criterion: for each set we compute the RankMe effective rank [30] which measures how many dimensions the representation effectively uses through the entropy of the normalized singular-value spectrum. Since the effective rank is bounded and biased by the number of rows, both sets are first reduced to the same number of samples *n* = min(*n*_train_, *n*_val_) by subsampling the larger one and averaging over 10 draws, and the resulting difference is normalized by the shared ceiling min(*n, d*), where *d* is the embedding dimension. This overfitting index stays close to zero as long as the encoder generalizes and rises once the training-graph representation becomes richer than what is reachable for unseen samples. We track the fold-averaged index together with its running minimum and declare the onset of overfitting at the first evaluation whose index exceeds that minimum by more than a tolerance of 0.05, provided the excess persists for three successive evaluations. The model is then retrained on the full training dataset for the average of the number of epochs at which this onset occurred during cross-validation (or, if no onset was detected, for the full epoch budget), yielding embeddings that generalize to unseen samples rather than overfitting the training data.

### 4.5 POME: implementation

POME is available as a Python package and is implemented using PyTorch [31] and PyTorch Geometric [32]. It is optimized for training on GPU, but is also applicable to smaller datasets on CPU. The required input format is based on pandas’ dataframe class and is described in detail on POME’s GitHub repository.

### 4.6 Competitor methods

Data imputation with MissForest [22] was carried out with the MissForest Python package [33], which natively supports both numeric and categorical variables. Auto-Complete [14] was run using the implementation provided in the corresponding GitHub repository (https://github.com/sriramlab/AutoComplete). For *k*-NN imputation, we used scikit-learn’s class `sklearn.impute.KNNImputer`. Since both *k*-NN imputation and AutoComplete are restricted to binary (i.e. two-class) categorical data, we employed one-hot-encoding on for non-binary categorical variables. In some cases, AutoComplete left a small amount of missing data points unimputed. When computing multiclass accuracy to assess imputation performance for categorical variables, we treated the unimputed values as a separate “NA” class. To compute mean absolute errors for imputation results on numeric variables, we set the error for the unimputed values to the maximal error for successfully imputed values on the respective dataset.

For applying UMAP [9] to POM data, we fitted two separate UMAP models on numeric and categorical variables separately and then used the intersection operator to combine both models into one, as suggested in the UMAP documentation [10]. Since UMAP does not intrinsically support distance metrics capable of handling missing data, we added two missingness-aware distance measures ourselves: for the numeric UMAP model, we implemented a pairwise removal-based Euclidean distance function, which scales distances by the inverse ratio of the number of present values in both input vectors. This implementation directly corresponds to the missingness-aware Euclidean distance implemented in scikit-learn’s `sklearn.metrics.pairwise.nan_euclidean distances` [34]. Analogously, we added a pairwise removal-based Hamming distance function, which is used for embedding categorical variables. In addition, we encountered that when passing a dataset with missingness (and hence setting the input parameter `ensure_all_finite=“allow-nan”`) and with more than 4096 input samples, the Python package `pynndescent` used in UMAP for the approximate computation of the underlying *k*-NN graph reports several issues due to missing data in the input. We fixed this issues by changing the UMAP implementation to always exactly compute the underlying *k*-NN graph.

In order to be able to compare against UMAP in the linear probing experiment, we further extended UMAP’s inductive capability (i. e., the capability to embed unseen samples into an already fitted embedding model via a call to `transform()`): First, since the missingness-aware distances above were only used when fitting the model, we added bipartite (new samples × training samples) variants of both measures and modified `transform()` to compute the new-to-training distances exactly with these measures whenever `ensure_all_finite=“allow-nan”` is set, again bypassing `pynndescent`’s approximate *k*-NN index. Second, UMAP models combined through the intersection operator do not support `transform()` at all, as they retain only the combined fuzzy simplicial set and the joint training embedding, but none of the fitted state of their components. We therefore implemented a function `transform combined()` which, given the fitted per-modality models, the combined model, and the new data per modality, (i) computes one bipartite fuzzy simplicial set per modality using that modality’s own distance measure, (ii) combines them with the same intersection operation used for the training fuzzy simplicial sets, (iii) initialises each new sample at the weighted mean position of its training neighbours, and (iv) optimises the new positions against the fixed combined training embedding, thereby placing the new samples in the joint embedding space. This requires that all component models were fitted on the same samples in the same order and that both the component and the combined models are retained after fitting.

### 4.7 Datasets

We evaluated POME on three real-world biomedical POM datasets: TCGA-LUAD, MIMIC-IV, and HANCOCK. We here provide brief descriptions of these datasets, overview tables are provided in the supplement (Tables S1–S2).

HANCOCK [17] is a multimodal datasets that comprises monocentric, real-world data of 763 head and neck cancer patients. We downloaded the different modules storing ICD-10 codes, as well as demographic annotations, blood data, pathological data, and target information from the corresponding GitHub repository and aggregated them into a joint representation. We chose target variables that were already designated as such in the dataset’s GitHub repository.

For the TCGA-LUAD dataset, we retrieved phenotype data for all lung adenocarcinoma patients from a standardized and cleaned version of the TCGA Pan-Cancer cohort [16]. After extraction of only lung adenocarcinoma patients, we removed fully missing columns and columns with only one unique value. Furthermore, we manually enriched the dataset with three more phenotypes from the exposure module of the TCGA-LUAD raw phenotype data accessed via the UCSC Xena Browser [35]: `cigarettes_per_day`, `pack_years_smoked`, and `years_smoked`. The final dataset encompasses 566 patients and 76 variables in total, with three held-out target variables for linear probing. Target variables were chosen as common patient outcome features, similar to those used in the HANCOCK dataset.

MIMIC-IV is a publicly available dataset containing de-identified information on patients admitted to Beth Israel Deaconess Medical Center between 2008 and 2019 [18]. We used the neutropenic fever and aplasia cohorts curated by Rahimi et al. [19], each comprises cancer patients treated with chemotherapy and includes a binary target variable indicating whether the corresponding chemotherapy side effect occurred during 45-day (aplasia) or 30-day (neutropenic fever) observation windows after discharge. As for HANCOCK, we chose these two target variables, as they were already used as prediction targets in the original publication [19]. To enable a joint analysis, we constructed a unified cohort by selecting only admissions present in both cohorts such that the target variables for both conditions were available for each admission. We included 100 numeric variables as input features, corresponding to the most frequently measured laboratory variables in the cancer chemotherapy cohort, averaged over the last 14 days prior to discharge.

### 4.8 Details on the unsupervised sample stratification and visualization experiments

For the comparison of unsupervised sample stratification between POME and UMAP embeddings, we applied Leiden clustering [23] on top of a *k*-NN graph. We used the Python implementation of Leiden clustering from [36]. For the *k*-NN graph computation, we set *k* to 5% of the total number of samples in the respective dataset, used the Euclidean distance for UMAP embeddings, and the squared Euclidean distance for POME embeddings (see justification above). For analyzing clusterability of POME’s sample embeddings in two dimensions, we computed Silhouette scores [37] and Davies-Bouldin indices [38] with respect to standard Euclidean distance, as visual perception is usually best quantified by Euclidean distance. The differential one-versus-all cluster analysis was performed using the missingness-aware implementations of the Mann-Whitney U test (for numeric variables) and the *χ*^2^-test (for categorical variables) from the NApy Python package [39]. The survival analysis on HANCOCK clusters 0 and 4 directly follows the survival analysis from the original HANCOCK publication [17] and is based on the lifelines Python package [40].

### 4.9 Details on the adjuvant therapy recommendation case study

All patients represented in the HANCOCK dataset underwent surgery [17]. For our AT recommendation use case, we trained POME using exclusively those variables whose values were determined at the time of or prior to surgery, i. e., we excluded all targets along with their associated variables (Table S2). AT modality status was determined based on the entries of the Boolean variables `adjuvant_systemic_therapy` and `adjuvant_radiotherapy`. The variable `year_of_initial_diagnosis` was used to split the patients into query and reference patients.

### 4.10 Details on the linear probing experiments

For our linear probing experiments we used scikit-learn’s `sklearn.linear model.LogisticRegression` (with L2 penalty, liblinear solver, C = 1, max iter = 1000). For each dataset we drew 10 random 80/20 train/test splits, fit the encoder on the training split, transformed the held-out test samples with POME’s inductive `transform()` function, fit the logistic regression on the training embedding and scored it on the test embedding, repeating each split with 5 embedding seeds (i.e. 50 fits per dataset, target and embedding size). Since the MIMIC-IV dataset contains admission-level target information and sometimes several admissions for the same patient, the splits were drawn at patient level, with all admissions of a patient assigned to the same partition. While this can lead to slightly imbalanced admission-to-split assignments, it prevents admissions from the same patient ending up in both the training and the test partition, which would constitute data leakage [41]. Since some target variables are not fully observed in the TCGA-LUAD dataset, samples with missing target values were excluded before fitting and scoring.

### 4.11 Details on the exploratory analysis of variable semantics

For clustering variable encoding vectors, we again used Leiden clustering on a *k*-NN graph based on pairwise cosine similarities and with *k* set to 5% of the number of variables in the MIMIC-IV dataset.

## Data availability

- The whole TCGA Pan-Cancer clinical dataset was downloaded from https://github.com/GerkeLab/TCGAclinical?tab=readme-ov-file. Additional clinical data for the TCGA-LUAD dataset was downloaded from https://xenabrowser.net/datapages/?cohort=GDC%20TCGA%20Lung%20Adenocarcinoma%20(LUAD)&removeHub=https%3A%2F%2Fxena.treehouse.gi.ucsc.edu%3A443.
- The HANCOCK dataset was downloaded from https://github.com/ankilab/HANCOCK_MultimodalDataset. The data for the variables `adjuvant_systemic_therapy`, `adjuvant_radiotherapy`, and `year_of_initial diagnosis` used for the AT modality recommendation use case was downloaded from the project website (https://www.hancock.research.uni-erlangen.org/download), as the GitHub repository does not contain these data.
- The MIMIC-IV dataset was accessed under a data use agreement through the PhysioNet project. Access to this dataset requires prior approval and completion of the relevant data use agreement and can be requested here: https://physionet.org/content/mimiciv/3.1/.

## Code availability

POME is available as Python package on GitHub (https://github.com/bionetslab/POME) and on PyPI (https://pypi.org/project/pome-py). Source code to reproduce the results reported in this article is available on GitHub (https://github.com/bionetslab/POME_evaluation).

## Author contributions

F.W. and D.B.B. conceived and designed the study and drafted the manuscript. F.W. implemented POME and the evaluation experiments. F.R and C.S. conducted the preprocessing of the MIMIC-IV dataset. L.A. assisted with the customization of UMAP for POM data. A.M.K. suggested the AT modality use case on the HANCOCK dataset. D.B.B. implemented the AT modality use case. D.B.B. and M.L. obtained funding and supervised the project. All authors provided critical feedback and discussions and assisted in interpreting the results and writing the manuscript.

## Ethics approval and consent to participate

Not applicable (only public data were used for this study).

## Consent for publication

Not applicable (only public data were used for this study).

## Supplementary information

The supplement contains eight supplementary figures (Figs. S1–S8) and two supplementary tables (Tables S1, S2).

