## Supplement for "POME: Graph-based embeddings for partially observed mixed-type data"

Fabian Woller, Lis Arend, Andreas M. Kist, Markus List,
Farnaz Rahimi, Christel Sirocchi, David B. Blumenthal

#### Supplementary figures

S1 Results of imputation experiments with alternative binning strategies. 2
S2 Results of imputation experiments with embedding size $d = 16$. . . . . 3
S3 Results of imputation experiments with embedding size $d = 32$. . . . . 4
S4 Results of unsupervised visualization experiments separated by embedding size. . . . . 5
S5 Number of POME training epochs in linear probing analysis . . . . . . 6
S6 Effect of mixed-type data integration on linear probing. . . . . . . . . 7
S7 Runtime and memory benchmark of POME. . . . . . . . . . . . . . . . . 8
S8 Imputation results for increasing numbers of training epochs . . . . . . 9

#### Supplementary tables

S1 Dataset overview. . . . . . . . . . . . . . . . . . . . . . . . . . . . . 10
S2 Overview of target variables. . . . . . . . . . . . . . . . . . . . . . . . 10

### 1 Supplementary figures

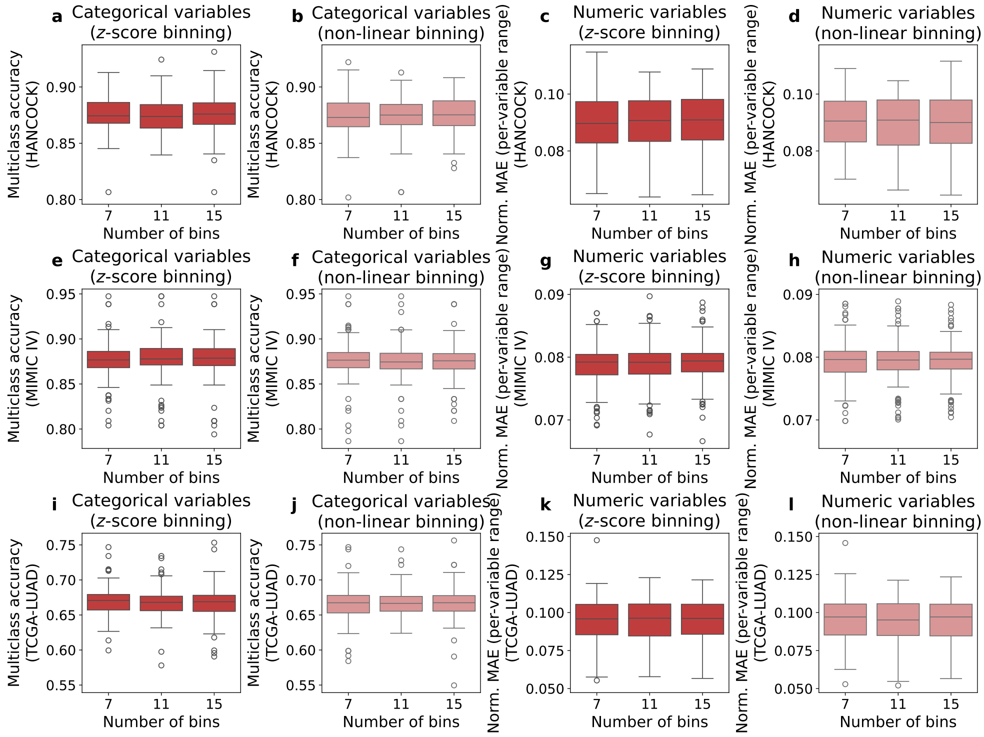


**Fig. S1** Results of imputation experiments with alternative binning strategies. Rows correspond to our three analyzed datasets HANCOCK (**a**–**d**), MIMIC-IV (**e**–**h**), and TCGA-LUAD (**i**–**l**). The first two columns compare results of $z$-score binning and non-linear binning on categorical variables (i.e. multiclass accuracies). The third and fourth columns compare the effect of $z$-score binning and non-linear binning on numeric variables (measured by normalized mean absolute errors). In all plots, errors and accuracies are aggregated over all simulated missingness ratios and are shown for varying numbers of discretization bins on the $x$-axis. The non-linear binning function $b_j : \mathcal{D}_j \to \{1, \ldots, K\}$ for a numeric variable $j$ is defined as follows: Let $x_{\min}$ and $x_{\max}$ be the empirical bounds of the data in $\mathcal{D}_j$ and $f : \mathbb{R} \to \mathbb{R}, f(x) = \text{sgn}(x) \cdot x^2$ be a quadratic transformation of input values. We first determine the boundary points in the transformed space by creating $B$ equidistant values $\tilde{e}_b = f(x_{\min}) + (b-1) \cdot \frac{f(x_{\max}) - f(x_{\min})}{B}$ for $b \in \{1, \ldots, B\}$. The resulting bin edges $e_b$ in the original domain $\mathcal{D}_j$ are recovered by inverting the quadratic transformation $f^{-1}(y) = \text{sgn}(y) \cdot \sqrt{|y|}$, i.e., $e_b = \text{sgn}(\tilde{e}_b) \cdot \sqrt{\tilde{e}_b}$. Based on these, the actual binning function $b_j(x)$ is defined as $b_j(x) = \max\{b \in \{1, \ldots, B\} \mid x \geq e_b\}$.

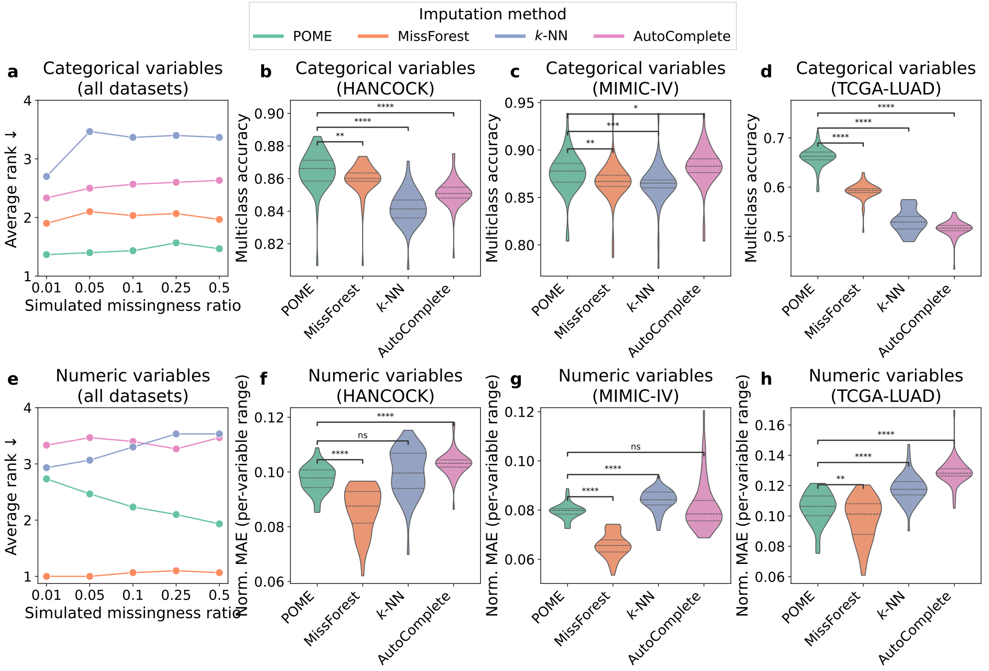


**Fig. S2** Results of imputation experiments with embedding size $d = 16$. **a** Average ranks of the compared methods on the categorical variables, jointly computed over the HANCOCK, MIMIC-IV, and TCGA-LUAD datasets (lower is better). **b–d** Multiclass accuracy scores on the categorical variables for each dataset separately, with violin plots visualizing results for all five simulated missingness ratios. **e** Average ranks of the compared methods on the numeric variables, jointly computed over the HANCOCK, MIMIC-IV, and TCGA-LUAD datasets (lower is better). **f–h** Mean absolute error on the numeric variables, normalized per variable by its min–max range, for each dataset separately, with violin plots visualizing results for all five simulated missingness ratios.

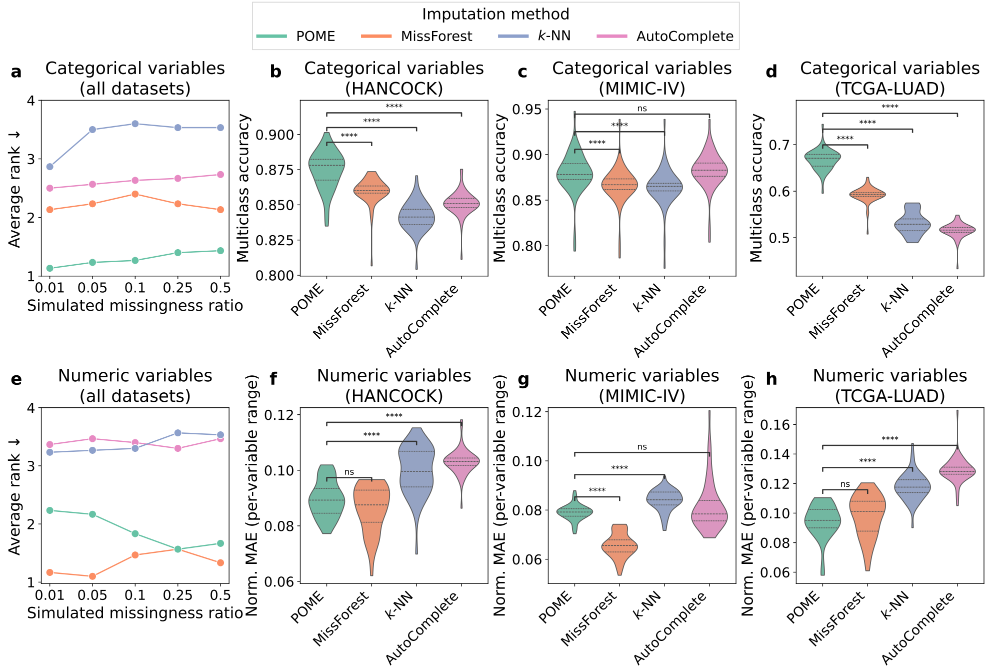


**Fig. S3** Results of imputation experiments with embedding size $d = 32$. **a** Average ranks of the compared methods on the categorical variables, jointly computed over the HANCOCK, MIMIC-IV, and TCGA-LUAD datasets (lower is better). **b–d** Multiclass accuracy scores on the categorical variables for each dataset separately, with violin plots visualizing results for all five simulated missingness ratios. **e** Average ranks of the compared methods on the numeric variables, jointly computed over the HANCOCK, MIMIC-IV, and TCGA-LUAD datasets (lower is better). **f–h** Mean absolute error on the numeric variables, normalized per variable by its min–max range, for each dataset separately, with violin plots visualizing results for all five simulated missingness ratios.

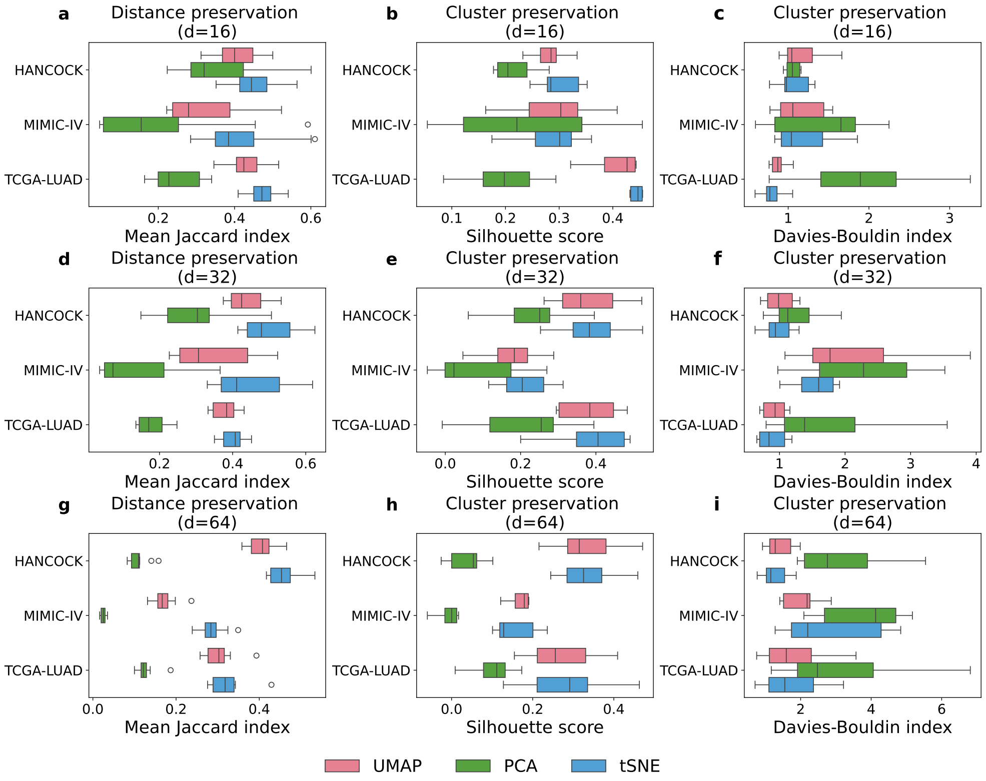


**Fig. S4** Results of unsupervised visualization experiments separated by embedding size. **a–c** Results for embedding size $d = 16$. **d–f** Results for embedding size $d = 32$. **g–i** Results for embedding size $d = 64$. First column: Local neighborhood preservation between POME embeddings and 2D projections. Second and third columns: 2D preservation of clusterings computed in $d$-dimensional embedding spaces via Leiden clustering, quantified via silhouette scores (second column) and Davies-Bouldin indices (third column) computed in 2D space.

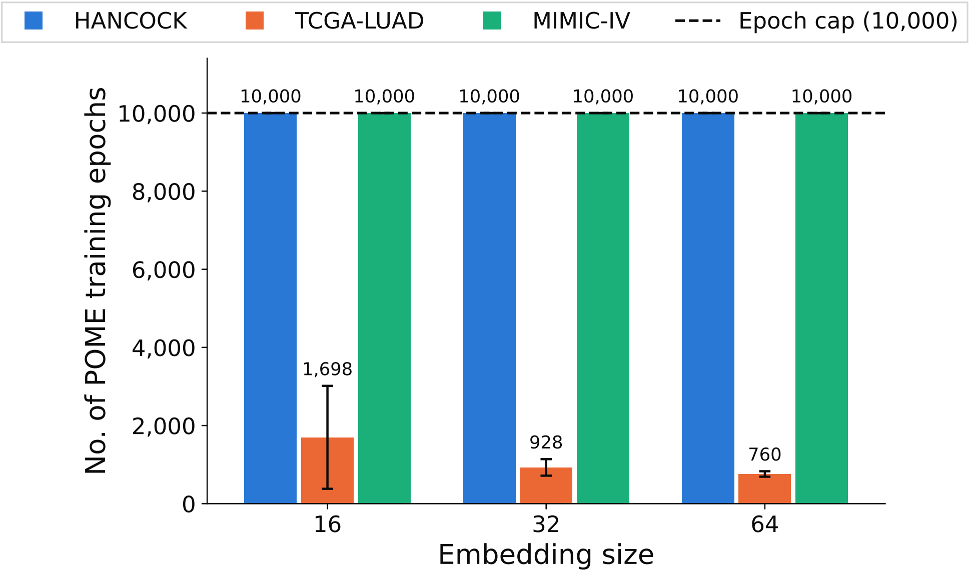


**Fig. S5** Number of training epochs used for linear probing computed by POME's unsupervised geometric criterion of the embedding matrix in order to avoid overfitting in the inductive mode. The bars show the mean and standard deviation over the 50 combined values of ten splits and five random seeds. For the TCGA-LUAD dataset, the resulting average number of training epochs is always below 2000 across all embedding sizes. On HANCOCK and MIMIC-IV, the suggested number of training epochs equals the manually set epoch limit of 10,000, which indicates that inductive embeddings of these two datasets benefit from longer training.

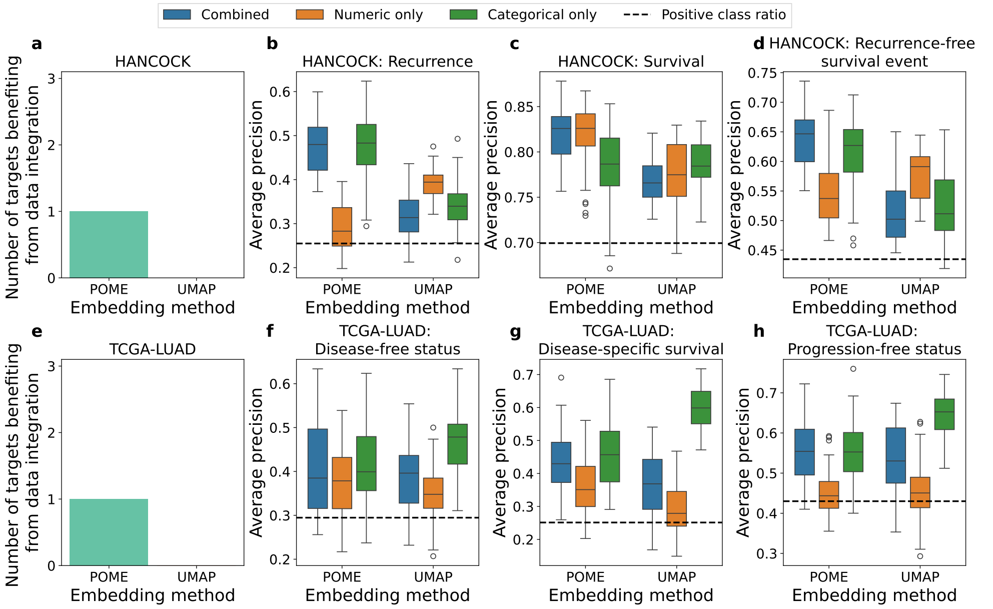


**Fig. S6** Effect of mixed-type data integration on linear probing. **a** Numbers of HANCOCK target variables which benefited from combining numeric and categorical variables, i. e., with the median average precision of embeddings computed on all variables ("Combined") being higher than both the median average precision of those based on only numeric variables ("Numeric only") and only categorical variables ("Categorical only"). **b–d** Results for binary HANCOCK target variables when using all variables from the dataset for embedding computation ("Combined"), only numeric variables ("Numeric only"), and only categorical variables ("Categorical only"). **e** Numbers of TCGA-LUAD target variables which benefited from combining numeric and categorical variables. **f–h** Results for binary TCGA-LUAD target variables, again separated by data types. All box plots show AP scores from the same train-test splits as used in the main linear probing analysis, obtained with five embeddings per embedding size which were generated with different random seeds (that is, each box plot summarizes $10 \cdot 5 = 50$ values). All shown results are based on an embedding size of $d = 64$, since larger embedding sizes tended to perform better on linear probing. MIMIC-IV is not included in this analysis, since the dataset only consists of numeric variables when excluding the two binary target variables.

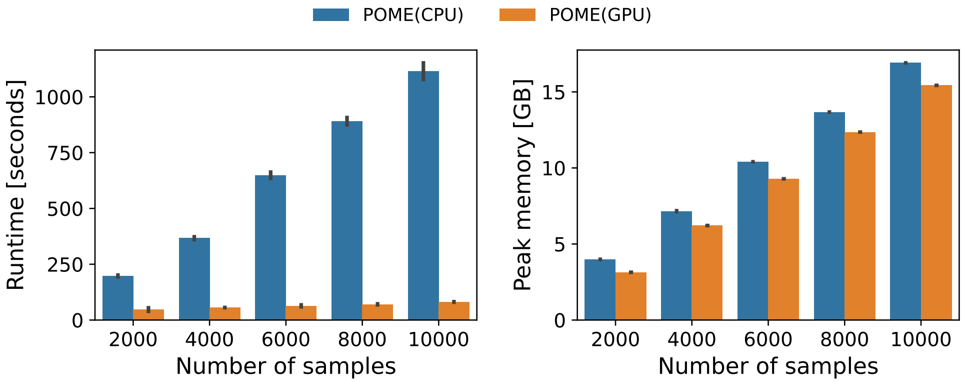


**Fig. S7** Runtime and memory benchmark of CPU- and GPU-based version of POME. Both versions were benchmarked on ten simulated datasets consisting of 100 features (50 numeric, 50 categorical) and increasing numbers of samples. For each variable, we randomly added 10% of missing values. Peak RSS memory of the CPU-based version was measured using the Python profiling tool `psutil`. For the GPU-based version, we measured peak VRAM using `torch.cuda.max_memory_allocated`. The tests were run on an Intel Xeon(R) Gold 6242R CPU @ 3.10GHz, and a NVIDIA RTX 6000 Ada Generation GPU, respectively. The bar plots show mean runtimes and memory requirements over five runs, with error bars visualizing standard deviation.

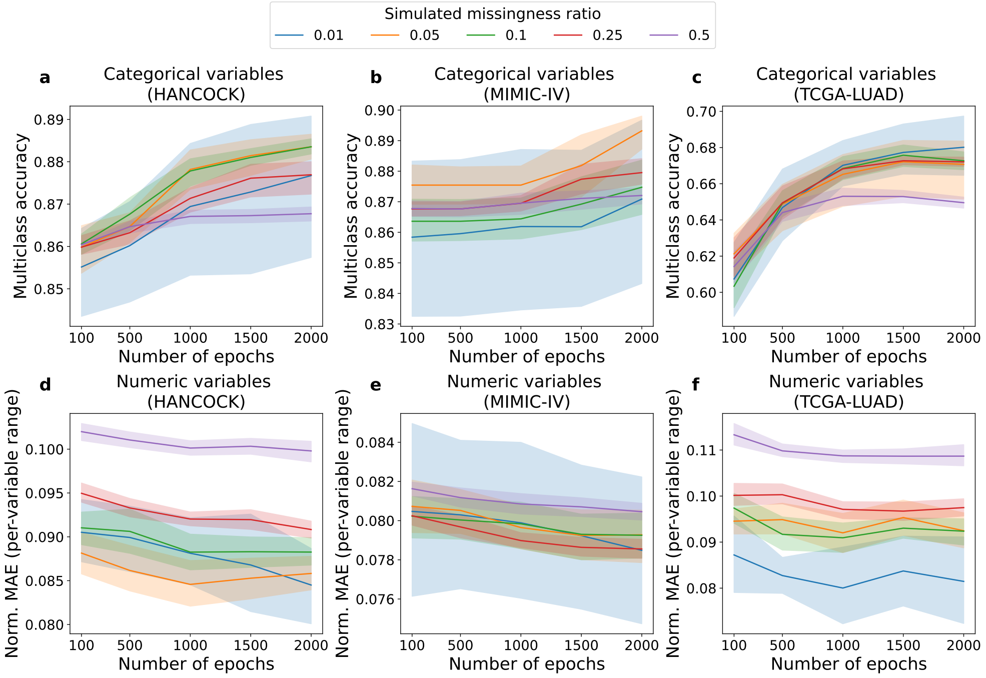


**Fig. S8** Imputation results for increasing numbers of training epochs with fixed embedding size $d = 32$. **a**–**c** Multiclass accuracy scores on the categorical variables for each dataset separately, with the number of steps that POME embeddings have been trained on the $x$-axis, and lines being colored by the five simulated missingness ratios. **d**–**f** Mean absolute error scores on the numeric variables for each dataset separately, with the number of epochs that POME embeddings have been trained on the $x$-axis, and lines being colored by the five simulated missingness ratios.

### 2 Supplementary tables

**Table S1** Dataset overview. The missingness ratio is defined as $|\mathcal{O}|/(|\mathcal{S}| \cdot |\mathcal{V}|)$, where $\mathcal{S}$, $\mathcal{V}$, and $\mathcal{O}$ are the sets of all samples, variables, and observed sample-variable pairs.

| | | | Variables | | | |
|---|---|---|---|---|---|---|
| Dataset | $\lvert\mathcal{S}\rvert$ | Missingness ratio | $\lvert\mathcal{V}\rvert$ | $\lvert\mathcal{N}\rvert$ | # Targets | #Target-associated |
| HANCOCK | 763 | 7.1% | 86 | 26 | 3 | 4 |
| MIMIC-IV | 4428 | 53.1% | 102 | 100 | 2 | 0 |
| TCGA-LUAD | 566 | 19.7% | 84 | 14 | 3 | 5 |

**Table S2** Overview of target variables. Target and target-associated variables were excluded when computing the POME embeddings used for the unsupervised sample stratification experiments, for the linear probing experiments, and for the adjuvant therapy modality recommendation study.

| Dataset | Target | Positive class ratio | Associated variables |
|---|---|---|---|
| HANCOCK | RFS event | 0.397 | Days to RFS event |
| HANCOCK | Survival status | 0.721 | Survival status with cause, Days to last information |
| HANCOCK | Recurrence | 0.232 | Days to recurrence |
| MIMIC-IV | Aplasia | 0.212 | None |
| MIMIC-IV | Neutropenic fever | 0.051 | None |
| TCGA-LUAD | Disease-free status | 0.397 | Disease-free (months) |
| TCGA-LUAD | Disease-specific survival status | 0.721 | Months of disease-specific survival, Overall survival status, Overall survival (months) |
| TCGA-LUAD | Progression-free status | 0.232 | Progress-free survival (months) |
